# Depletion of Chloroplast HSP70B Triggers Proteostasis Collapse and Compromises Thylakoid Membrane Integrity in *Chlamydomonas*

**DOI:** 10.64898/2026.04.02.716084

**Authors:** Anna Probst, Stefan Schmollinger, Jonas Berg, Ann-Katrin Unger, Daniela Strenkert, Stefan Geimer, Frederik Sommer, Michael Schroda

## Abstract

Chloroplast HSP70 is an essential component of the plastid proteostasis network, supporting protein folding, complex assembly and disassembly, and stress acclimation. Despite extensive genetic evidence for its essentiality, the cellular consequences of reduced chloroplast HSP70 activity remain poorly defined. Here, we investigated the function of the sole chloroplast HSP70 in *Chlamydomonas reinhardtii*, HSP70B, using an inducible artificial microRNA approach that reduced HSP70B abundance to below 30% of wild-type levels. HSP70B depletion resulted in cell division arrest and extensive proteome remodeling, characterized by strong upregulation of proteins involved in chloroplast protein quality control and membrane remodeling. Notably, this response was accompanied by increased abundance of protein quality control components in the endoplasmic reticulum, cytosol, and mitochondria, indicating pronounced proteostasis cross-talk between cellular compartments. In contrast, chloroplast and cytosolic ribosomes, photosynthetic and respiratory protein complexes, and central metabolic enzymes were broadly depleted, consistent with a collapse of cellular proteostasis. At the ultrastructural level, HSP70B-depleted cells exhibited lesions at thylakoid membrane conversion zones previously described in VIPP1-depleted cells. Accordingly, higher-order oligomeric forms of VIPP1 accumulated, and cells displayed extreme sensitivity to high-light stress. These findings confirm HSP70B as a key regulator of VIPP1 oligomer dynamics and highlight its central role in coordinating chloroplast membrane remodeling with cellular proteostasis in *Chlamydomonas*.

**One-sentence summary:** Depletion of chloroplast HSP70B causes cell division arrest, proteostasis collapse, impaired VIPP1 oligomer dynamics with aberrant thylakoid structures, and increased light sensitivity.

## Introduction

Chaperones of the HSP70 class are key regulators of cellular proteostasis (Balchin et al., 2016). They assist in the folding of newly synthesized proteins, the refolding of misfolded or aggregated proteins, the translocation of proteins across membranes, and the assembly, disassembly, or remodeling of protein complexes (Rosenzweig et al., 2019).

HSP70s consist of an N-terminal ∼45-kDa nucleotide-binding domain (NBD), a ∼15-kDa β-sandwich substrate-binding domain (SBDβ), a ∼10-kDa α-helical lid domain (SBDα), and a disordered C-terminal tail (Mayer and Bukau, 2005). Their activity relies on an ATP-driven conformational cycle that is tightly regulated by specific co-chaperones, most notably J-domain proteins and nucleotide exchange factors (NEFs) (Kityk et al., 2015; Rosenzweig et al., 2019). In the ATP-bound state, SBDβ and SBDα dock onto the NBD, which suppresses ATP hydrolysis and allows the transient binding of protein substrates—typically exposing hydrophobic stretches flanked by basic residues—to the substrate-binding pocket (Rüdiger et al., 1997). Substrates are commonly delivered by J-domain co-chaperones, which also confer substrate specificity (Craig et al., 2006). Substrate binding and interaction of the J domain with the NBD stabilize a conformation competent for ATP hydrolysis, leading to dissociation of SBDβ and SBDα from the NBD. Upon ATP hydrolysis, the SBDα lid closes over the substrate-binding pocket of SBDβ, thereby preventing substrate dissociation. In the ADP-bound state, the NBD and SBD are largely independent and connected by an interdomain linker. Release of ADP and rebinding of ATP, which reopens the nucleotide-binding cleft and promotes substrate release, is facilitated by NEFs (Bracher and Verghese, 2015).

HSP70s are constitutively present in the cytosol/nucleus, endoplasmic reticulum, mitochondria, and chloroplasts and often are increasingly expressed in response to various proteotoxic stress conditions (Schroda and deVitry, 2023). Whereas the unicellular green alga *Chlamydomonas reinhardtii* (Chlamydomonas) harbors a single chloroplast HSP70 (HSP70B), two chloroplast HSP70s are present in *Arabidopsis thaliana* (Arabidopsis; cpHsc70-1 and cpHsc70-2), and three in the moss *Physcomitrium patens* (Hsp70-1 to -3) (Trösch et al., 2015). Chlamydomonas stromal HSP70B cooperates with six J-domain proteins belonging to four distinct phylogenetic clades, represented by CDJ1, CDJ2, CDJ3–5, and CDJ6. Arabidopsis employs 26 J-domain proteins in the chloroplast, only 19 of which belong to the four conserved clades, indicating both an expansion of HSP70 function and further specialization towards clients in land plants (Chiu et al., 2013; Trösch et al., 2015). Chlamydomonas, moss, and Arabidopsis each possess two chloroplast NEFs termed chloroplast GrpE homologs (CGE1 and CGE2) (Trösch et al., 2015).

A role for Chlamydomonas HSP70B in chloroplast protein quality control is supported by its ability to prevent aggregation of denatured malate dehydrogenase and to assist in the refolding of denatured luciferase in vitro, with efficient refolding requiring the presence of CDJ1 and CGE1 (Veyel et al., 2014). HSP70B, CDJ1, and CGE1 were found in a complex with chloroplast HSP90C, possibly forming a chloroplast ‘foldosome’ (Willmund et al., 2008; Schroda and Mühlhaus, 2009). Interestingly, two J-domain proteins from the same clade as CDJ1 in Chlamydomonas, DJA5 (DjA24) and DJA6 (DjA26), were shown to bind ferrous iron and deliver it via their J domains to the SUFBC_2_D complex for iron–sulfur cluster biogenesis, independently of cpHsc70 activity (Zhang et al., 2021). In Arabidopsis, the J-domain protein J20 (DjC26), which belongs to a clade absent in Chlamydomonas, is likewise involved in chloroplast protein quality control by delivering inactive forms of deoxyxylulose 5-phosphate synthase (DXS), the key enzyme of the MEP pathway, to cpHsc70 for folding or degradation (Pulido et al., 2016).

A role for chloroplast HSP70 in protein complex (dis)assembly is exemplified by CDJ2/DjC73, which delivers the ESCRT-III family member VIPP1 to its chaperone partner. Chlamydomonas HSP70B was shown to assist in the assembly and disassembly of higher-order VIPP1 oligomers in vitro and in cell extracts (Liu et al., 2005; Liu et al., 2007). Similarly, Arabidopsis cpHsc70 promotes the disassembly of VIPP1 oligomers in vivo (Li et al., 2025). VIPP1 forms baskets composed of stacked rings (Gupta et al., 2021; Liu et al., 2021), helical rods (Liu et al., 2007; Theis et al., 2019; Gachie et al., 2025), as well as spirals, polygons, and carpets (Junglas et al., 2025; Naskar et al., 2025; Pan et al., 2025). These oligomeric structures remodel chloroplast membranes and are implicated in the biogenesis and maintenance of thylakoid membranes (Nordhues et al., 2012; Zhang et al., 2012).

A role for Arabidopsis cpHsc70 in chloroplast protein import was proposed based on the stable association of cpHsc70-1 and cpHsc70-2 with TOC–TIC components and impaired import of chloroplast protein precursors in young seedlings lacking one of the two Hsc70s (Su and Li, 2010). Support for an involvement of the HSP70 system in chloroplast protein import also came from studies in moss, where import was impaired in mutants with reduced chloroplast HSP70 or CGE activity (Shi and Theg, 2010). In contrast, HSP70B was not found to interact with the TOC–TIC supercomplex in Chlamydomonas (Ramundo et al., 2020), and Hsp70 was absent from cryo-EM structures of purified TOC–TIC complexes from Chlamydomonas and land plants. Instead, these structures provide strong evidence that the Ycf2–FtsHi complex constitutes the chloroplast protein import motor (Jin et al., 2022; Liu et al., 2023; Liang et al., 2024).

Whether related to protein quality control, VIPP1 (dis)assembly, protein import, or a yet unknown function, chloroplast HSP70 activity is essential, as evidenced by the embryo-lethal phenotypes of Arabidopsis cp*hsc70-1 cphsc70-2* double knock-outs as well as moss *hsp70-2* and *cge1 cge2* mutants (Su and Li, 2008; Shi and Theg, 2010). While *Arabidopsis cphsc70-2* knock-out plants show no obvious phenotype, *cphsc70-1* mutants exhibit variegated cotyledons, irregular leaf margins, lesions on true leaves, shortened roots, and reduced biomass (Su and Li, 2008; Latijnhouwers et al., 2010; Ding et al., 2022). These phenotypes are exacerbated by high temperature, drought, and osmotic stress (Latijnhouwers et al., 2010; Ding et al., 2022; Li et al., 2025). Plants expressing an artificial microRNA targeting both *cpHsc70s* are nearly albino, severely growth-retarded, display drastically reduced levels of photosystem subunits, and contain no or only small chloroplasts that are often swollen and lack organized thylakoids (Latijnhouwers et al., 2010). In Chlamydomonas, *HSP70B* antisense lines with only a mild reduction in HSP70B abundance show increased sensitivity of PSII to high-light exposure and delayed recovery, whereas the opposite phenotype is observed upon HSP70B overexpression (Schroda et al., 1999). Moreover, *HSP70B* antisense lines exhibit a weaker and delayed accumulation of heat shock proteins upon heat stress (Schmollinger et al., 2013).

Here, we used an inducible amiRNA approach to reduce chloroplast HSP70B levels to below 30% of wild-type abundance. Partial depletion of HSP70B caused cell division arrest and extensive proteome remodeling affecting protein quality control across compartments, ribosome abundance, photosynthetic and respiratory complexes, central metabolism, and thylakoid membrane integrity. These effects were accompanied by impaired VIPP1 oligomer dynamics and pronounced sensitivity to high-light stress.

## Results

### Downregulation of HSP70B results in growth arrest and increased cell diameter

To reduce the HSP70B content in Chlamydomonas cells, we employed an inducible amiRNA system based on the nitrate reductase (*NIT1*) promoter (Schmollinger et al., 2010). Here, the expression of an amiRNA that targets sequences in the third exon of the *HSP70B* mRNA is induced by switching from ammonium to nitrate as the sole nitrogen (N) source (Supplemental Figure S1A). We transformed the *HSP70B*-amiRNA construct pMS542 (*70B*-amiR) into the arginine-auxotrophic Chlamydomonas strain cw15-325 which, unlike most Chlamydomonas laboratory strains, contains intact *NIT1* and *NIT2* genes and can therefore utilize nitrate as an N source. Using the wild-type (WT) *ARG7* gene in the pMS542 vector, we selected for arginine-prototrophic transformants (depicted with #). In some of these transformants, 24 hours after switching from ammonium to nitrate, HSP70B protein contents were reduced to as little as 10% of the WT content (Figure 1A; Supplemental Figure S2A). With ammonium as N source, HSP70B protein levels were largely unchanged compared to a control line generated with the empty amiRNA vector pMS539.

**Figure 1.**
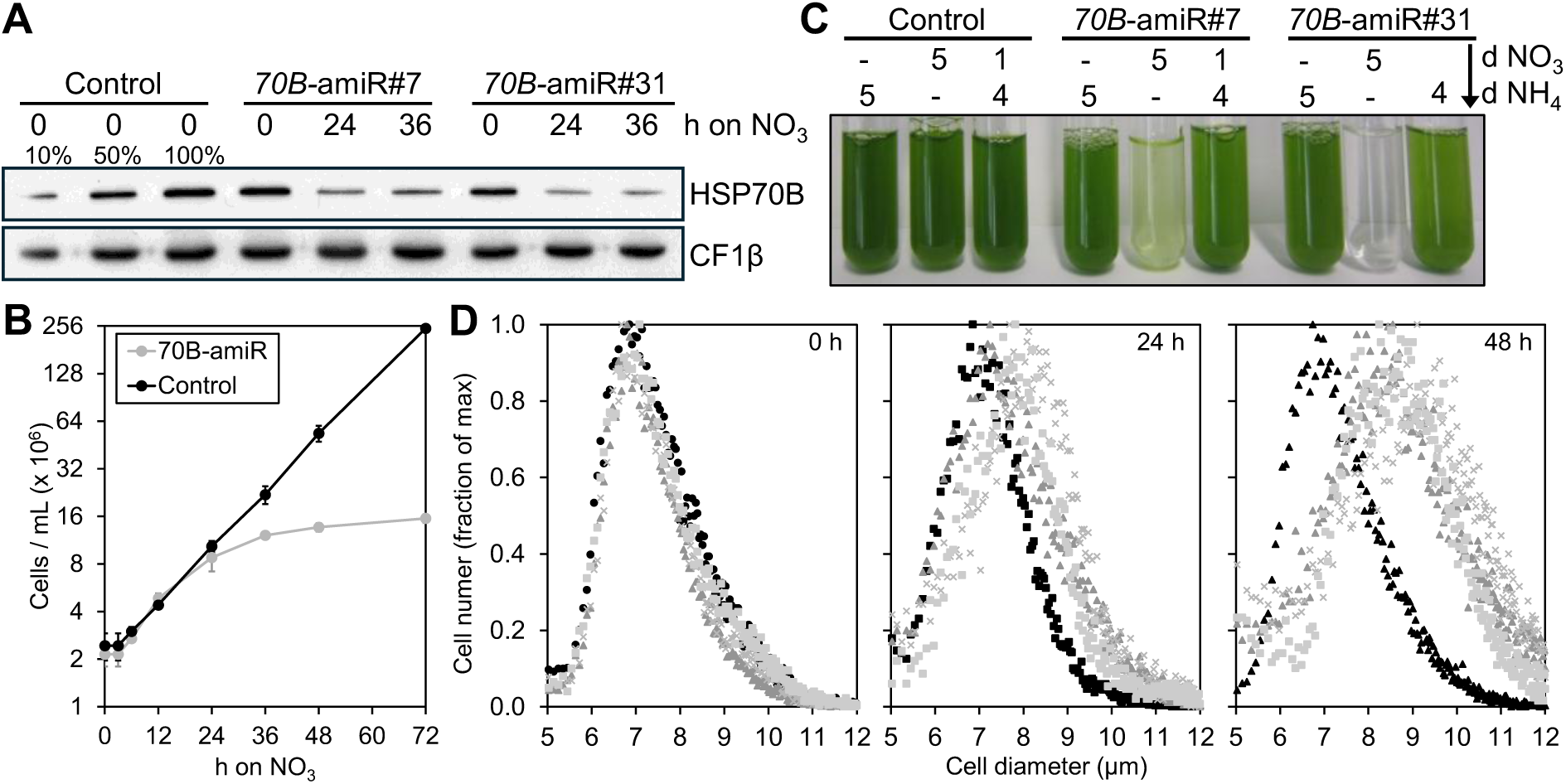
Analysis of cell growth and cell diameters in *70B*-amiR lines. **A)** Monitoring HSP70B protein abundance in two *70B*-amiR lines versus control. Total cell proteins of a control strain generated with the empty amiRNA vector pMS539 and two transformants generated with vector pMS542 (#7, #31) (Supplemental Figure 1A) were extracted before, and for the *70B*-amiR lines 24 h and 36 h after switching from ammonium to nitrate as N source (h on NO_3_). The content of HSP70B relative to the loading control CF1β was analyzed by immunoblot analysis of total proteins corresponding to 0.5 μg chlorophyll. **B)** Growth of three *70B*-amiR lines versus control. The cell number per mL culture of three control lines generated with pMS539 (black) and three *70B*-amiR lines (#7, #16, #31, gray) was determined at the indicated time points after switching from ammonium to nitrate (time point 0). The cultures were continuously diluted to 10^6^ cells/mL with fresh TAP-NO_3_ once a cell density of 5 x 10^6^ cells/mL was reached to keep the cells constant in mid-log phase. Mean values from the three control and three *70B*-amiR lines are shown. Error bars indicate SDs. **C)** Growth of two *70B*-amiR lines versus control. Lines were grown in medium with ammonium (d NH_4_) or nitrate (d NO_3_) for 5 days or transferred back to medium with ammonium for 4 days after 1 day in medium with nitrate. **D)** Cell size distribution of three *70B*-amiR lines versus control. The number of cells (fraction of max) with a given cell diameter (μm) is plotted against the cell diameter at the indicated time points after switching the N source in a control line (black) and three *70B*-amiR lines (gray squares (#2), triangles (#7), crosses (#15)).

For the first ∼3 h after switching the N source, we observed a delayed division rate in control and *70B*-amiR lines, after which both lines resumed exponential growth (Figure 1B). While the control lines kept growing exponentially, the division rate of the *70B*-amiR lines decreased ∼24 h after medium change and cells stopped dividing almost completely between 48 h and 72 h. *70B*-amiR lines grown on nitrate for 1 d resumed growth after transferring them back to ammonium for 4 d (Figure 1C). Changing the N source had no effect on the diameter distribution of dividing control line cells. However, this distribution shifted in favor of larger cells in the *70B*-amiR lines 24 hours after switching the N source, and this effect became more pronounced after 48 hours (Figure 1D). The *70B*-amiR lines in the cw15-325 strain background survived only for a few months on agar plates with ammonium as N source, presumably because of leaky expression of the *NIT1* promoter. After a laboratory move, we were no longer able to produce stable *70B*-amiR lines in the cw15-325 strain background.

To overcome this problem, we transformed the UVM4 strain (Neupert et al., 2009) that has defective *NIT1/2* genes with plasmids containing the wild-type *NIT1* and *NIT2* genes and selected transformants on nitrate-containing medium (UVM4-NIT). Furthermore, we constructed a new amiRNA vector (pMBS822) based on the Modular Cloning System (Crozet et al., 2018) to express the same amiRNA against *HSP70B* that was used in pMS542 (Supplemental Figure S1B). Some spectinomycin-resistant transformants obtained with pMBS822 showed the same inducible depletion of HSP70B after 24 h on nitrate-containing medium that was observed with cw15-325 transformants containing pMBS542. In contrast to lines in the cw15-325 background, the *70B*-amiR lines in the UVM4-NIT background were stable. A growth defect was observed in these lines, as well (Supplemental Figures S2B and S3).).

### HSP70B depletion increases protein quality control components and reduces photosynthetic light reaction complex subunits

To evaluate the molecular consequences of HSP70B depletion, we monitored the abundances of selected proteins by immunoblotting in *70B*-amiR lines of both strain backgrounds. As shown in Figure 2, the depletion of HSP70B resulted in substantial upregulation of chloroplast membrane remodelers VIPP1 and VIPP2 (Nordhues et al., 2012), stromal protease DEG1C (Theis et al., 2019), HSP70B co-chaperone CDJ1 (Willmund et al., 2008), chloroplast disaggregase CLPB3 (Kreis et al., 2023), chloroplast small heat shock proteins HSP22E/F (Rütgers et al., 2017), and autophagy marker ATG8 (Perez-Perez et al., 2010). Cellular contents of chloroplast HSP90C and cytosolic HSP90A chaperones (Schmollinger et al., 2013) increased also, but more mildly. Only 48 h after shifting the N source we observed a decline also of photosystem (PS) I core subunit PsaA as well of ATP synthase subunit CF1β in *70B*-amiR lines in the UVM4-NIT background.

**Figure 2.**
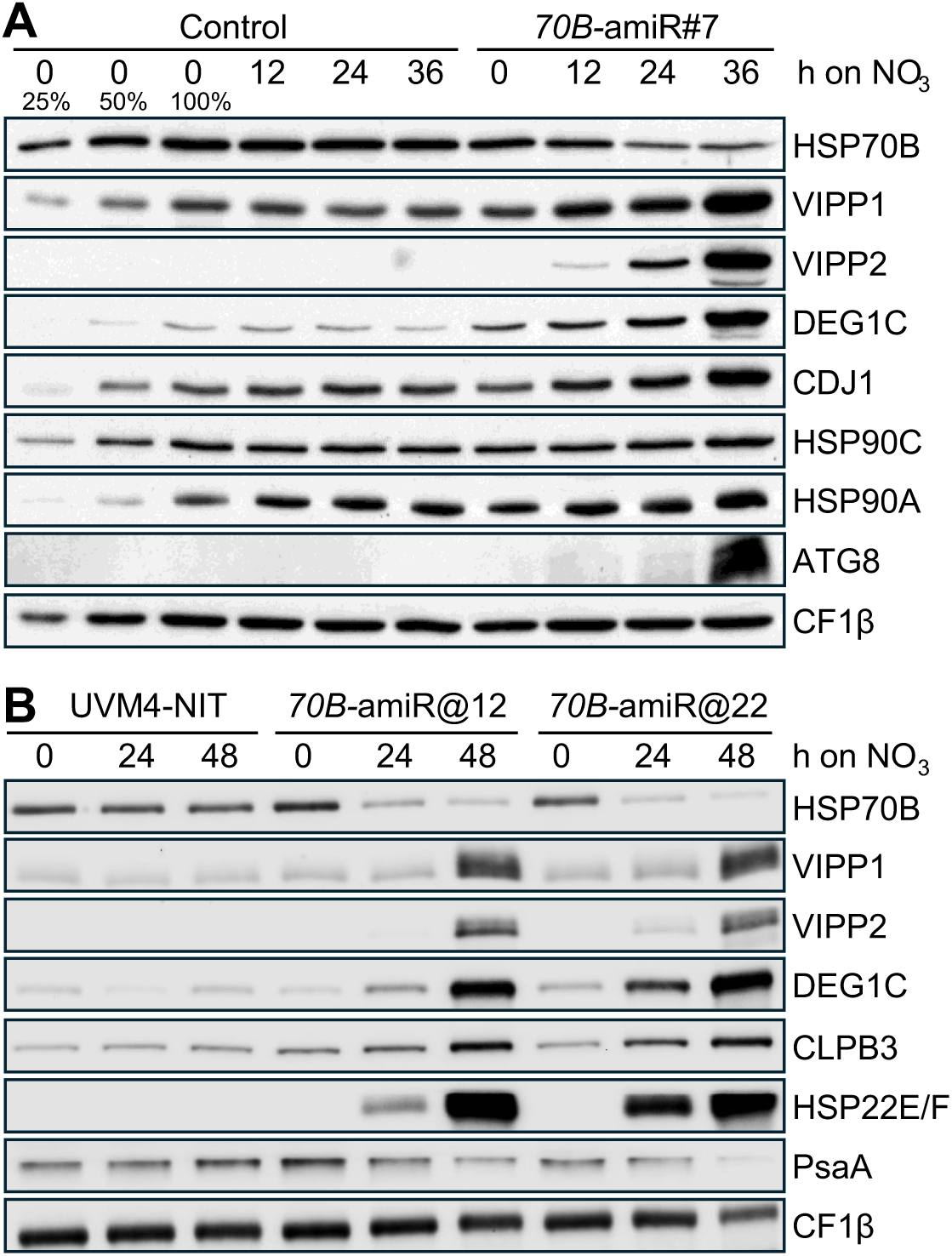
Immunoblot analysis of selected proteins in *70B*-amiR lines. **A)** Abundances of selected proteins in control and a *70B*-amiR line of the cw15-325 strain background (#). Total proteins before and after shifting cells from ammonium-to nitrate-containing medium to induce expression of the amiRNA construct were extracted and proteins corresponding to 0.5 to 2 µg chlorophyll were analyzed by immunoblotting. **B)** Abundances of selected proteins in control and a *70B*-amiR line of the UVM4-NIT strain background (@). (Protein analyses were done as in A).

### Changes in the cellular proteome upon shifting the N source are fully reversible in the UVM4-NIT line but not in the *70B*-amiR line

To assess the impact of HSP70B depletion on the cellular proteome in more detail, we performed label-free shotgun proteomics. We quantified 5983 proteins in the *70B*-amiR line @12 and its recipient strain UVM4-NIT1 before (0 h), 24 h and 48 h after shifting from ammonium to nitrate, and 24 h after shifting the N source back to ammonium (72 h, Supplemental Data Set S1). Principal component (PC) analysis showed that the four biological replicates for each time point and line clustered closely (Figure 3A). PC1 explained the largest proportion of the variance and primarily reflected the shift in N source. The similar positioning of the 24 h and 48 h nitrate time points along PC1 in both lines indicates that proteome adjustment to nitrate was largely complete within 24 h. However, the displacement along PC1 was more pronounced in the control than in the *70B*-amiR line, suggesting weaker proteome remodeling in the latter, possibly due to arrested cell division. Consistent with this interpretation, the control line readjusted its proteome more completely within 24 h after shifting back to ammonium, whereas readjustment was less complete in the *70B*-amiR line.

**Figure 3.**
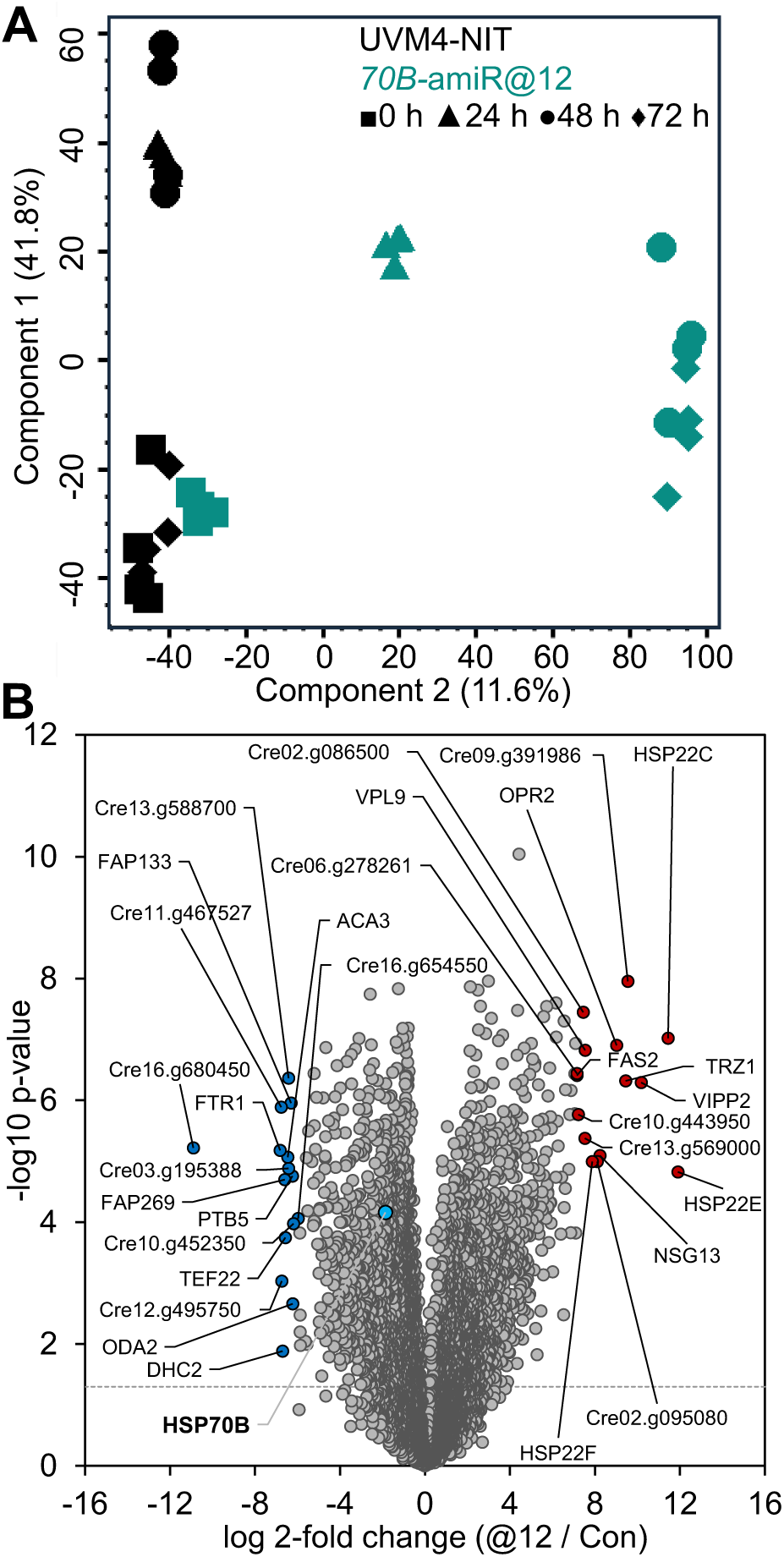
Overview of shotgun proteomics results. Cultures of the UVM4-NIT control and *70B-*amiR @12 lines were grown to mid-log phase on TAP-NH_4_, shifted at time point 0 h to TAP-NO_3_ for 48 h and then back to TAP-NH_4_ for another 24 h (72 h time point). **A)** Principle component analysis of the four biological replicates of each of the four time points analysed for each line. **B)** Volcano plot showing the abundances of 5983 proteins in *70B*-amiR *@*12 versus control 48 h after shifting the N source. Each data point represents the average of four replicates. The 15 proteins with highest fold-changes between *70B*-amiR*@*12 versus control are labeled (up – red, down – blue). HSP70B is shown for orientation.

PC2 separated the *70B*-amiR line from the control across all time points and is consistent with proteome changes associated with HSP70B depletion. The slight shift of the 0 h ammonium samples of the *70B*-amiR line along PC2 indicates that some proteome changes were already present under ammonium growth conditions, possibly due to leakiness of the *NIT1* promoter. The stable positioning of later time points along PC2 suggests that these HSP70B-associated proteome changes persisted throughout the experiment.

Presumably due to the leakiness of the *NIT1* promoter, the HSP70B content in the *70B*-amiR line was already reduced to ∼65% of wild-type levels in cells grown on ammonium (Figure 4). Upon switching to nitrate, the HSP70B content declined to 24% and 28% of WT levels after 24 h and 48 h, respectively, while the HSP70B content was only 40% of WT levels 24 h after shifting back to ammonium. Probably due to this reduced cellular HSP70B content before switching the N source, levels of the sensitive cpUPR markers VIPP2, HSP22E, and DEG1C were already 3-, 6.5-, and 2.5-fold higher in the *70B*-amiR line than in UVM4-NIT (Figure 4A-C), explaining the slight shift of 0 h ammonium samples of the *70B*-amiR line along PC2 in Figure 3A.

**Figure 4.**
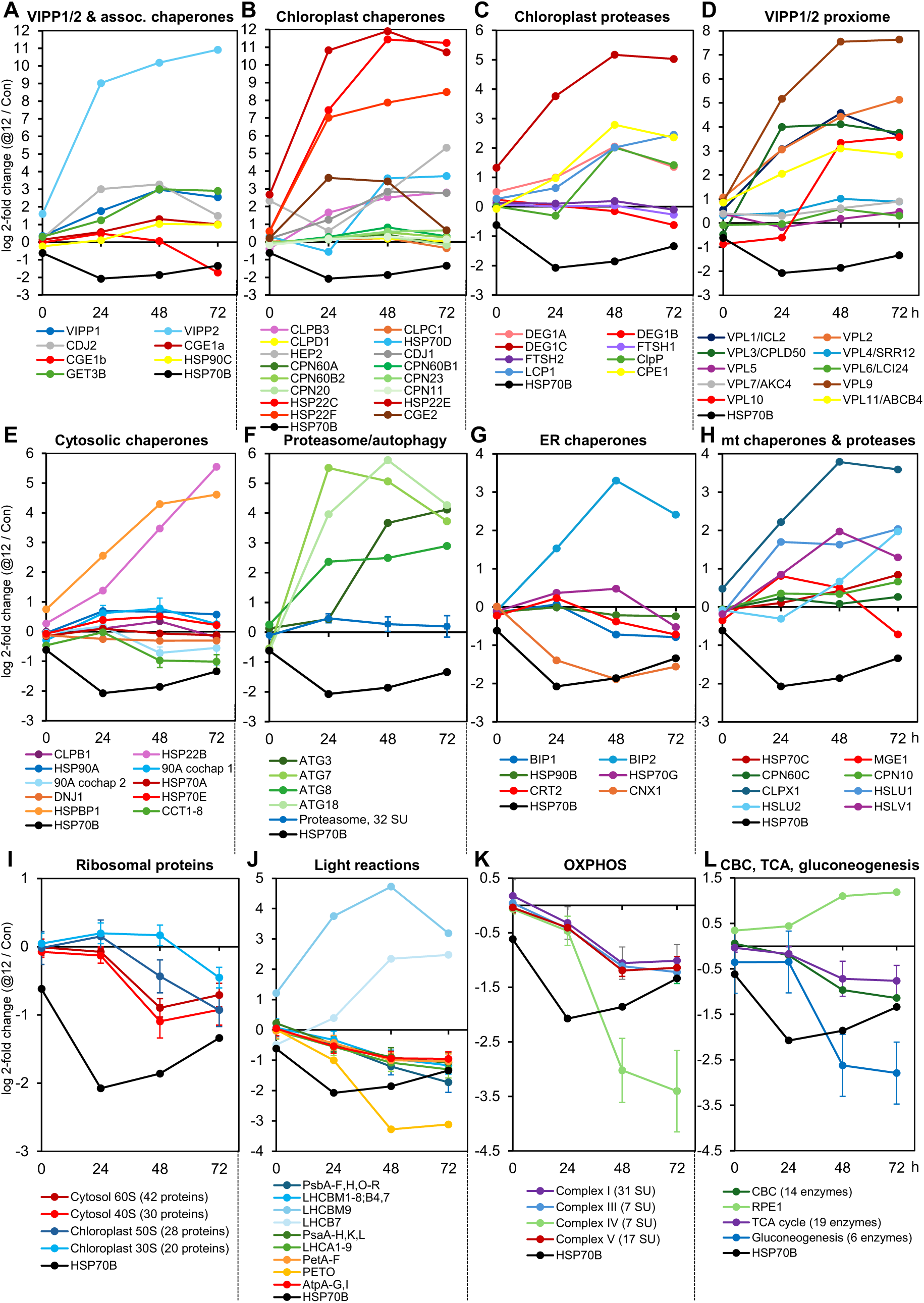
Monitoring cellular protein contents during HSP70B depletion and repletion by shotgun proteomics. Shown are log2-fold differences in the abundance of the indicated proteins between *70B-amiR* and control line. Values for each protein at each time point are the mean of 4 biological replicates. For more clarity, error bars indicating SDs are not shown for individual proteins but when multiple subunits (SU) of the same protein complex or enzymes of the same metabolic pathway were combined, as indicated in the legends. Values for HSP70B are shown in each panel to facilitate comparisons between the different Y-axis scales.

Among the 10 most upregulated proteins in the *70B*-amiR line 48 h after shifting the N source were chloroplast small heat shock proteins HSP22C and HSP22E/F and VIPP2 (Figures 3B and 4A, B), corroborating the immunoblot data of Figure 2. Furthermore, a phospholipase A1 (Cre09.g391986), a nuclear RNase Z-like protein (TRZ1), 12-oxophytodienoic acid reductase (OPR2), N-starvation expressed protein NSG13 (Abe et al., 2004), a major vault protein (Cre02.g095080), and the VIPP1-proximal protein VPL9 (Kreis et al., 2023) (Figures 3B and 4D). Among the 10 most downregulated proteins in the *70B*-amiR line compared to WT were the plasma membrane iron permease FTR1, P-type ATPase/cation transporter, plasma membrane ACA3, two cilia proteins, and six proteins of unknown function.

### HSP70B depletion strongly affects proteins involved in plastid membrane and protein homeostasis

Given the role of chloroplast HSP70 in plastid protein and membrane homeostasis (Schroda et al., 1999; Liu et al., 2007; Su and Li, 2008; Latijnhouwers et al., 2010; Schmollinger et al., 2013; Theis et al., 2020; Ding et al., 2022; Li et al., 2025), we compiled changes in the abundances of VIPP1/2 and associated proteins, cellular chaperones and proteases, ribosomal proteins as well as proteins involved in photosynthetic light reactions, respiration, and major metabolic pathways.

VIPP2 and VIPP1 themselves were strongly upregulated by >1100- and ∼8-fold, respectively, in the *70B*-amiR versus control line after 48 h of the N source shift (Figure 4A). Similarly, all (co-)chaperones found previously to be associated with VIPP1/2 were upregulated, including HSP70B co-chaperones CDJ2 and CGE1a (9.7- and 2.5-fold) (Liu et al., 2007), HSP90C (∼2-fold) (Heide et al., 2009) and GET3B (8-fold) (Kreis et al., 2023). In the group of other chloroplast chaperones, the small heat shock proteins HSP22C, E and F accumulated with similar kinetics to up to 3850-fold higher levels in the *70B*-amiR versus control line after shifting the N source (Figure 4B). HEP2, the escort protein of HSP70B (Willmund et al., 2008), HSP70B co-chaperones CDJ1 and CGE2 (Kreis et al., 2023), chloroplast disaggregase CLPB3, and truncated Hsp70 protein HSP70D accumulated in the *70B*-amiR line to 6- to 32-fold higher levels than in UVM4-NIT 48 h after shifting the N source (Figure 4B). In contrast, HSP100 members ClpC1 and CLPD1 and CPN60 chaperonin subunits (Zhao et al., 2019) were upregulated maximally 1.8-fold 48 h after the N source shift. Like chloroplast chaperones, chloroplast proteases also showed distinct response patterns to HSP70B depletion (Figure 4C): while stromal DEG1C was most upregulated (36-fold) in the *70B*-amiR line compared to UVM4-NIT, levels of FTSH1/2 (Malnoe et al., 2014) and DEG1B rather declined over time. Chloroplast processing enzyme CPE1 (Richter and Lamppa, 2003), lumenal DEG1A (Theis et al., 2019), plastid-encoded ClpP (Majeran et al., 2000), and Lon domain containing protein LCP1 (Shin et al., 2020) accumulated with similar kinetics to 4- to 7-fold higher levels in the *70B*-amiR line compared to UVM4-NIT.

The abundance of all ten VPL proteins found previously in the proximity of VIPP1/2 and upregulated under various chloroplast stress conditions (Kreis et al., 2023) increased in the *70B*-amiR versus control line (Figure 4D): most increased 48 h after the N source switch was VPL9 (∼200-fold), while VPL1, 2, 3, 10, 11 increased intermediately by ∼7- to 35-fold and VPL4-7 only mildly by at most 2-fold.

### HSP70B depletion has systemic consequences outside the chloroplast

Interestingly, the depletion of chloroplast HSP70B also resulted in specific changes in cytosolic chaperone abundances: levels of CLPB1 and HSP22B, involved in protein disaggregation (Queitsch et al., 2000), increased 1.3- and 11-fold, respectively, in the *70B*-amiR versus control line 48 h after the N source shift (Figure 4E). The abundance of HSP70A in the *70B*-amiR line did not change, while levels of its co-chaperone DNJ1, involved in protein folding (Silflow et al., 2011), declined by 20% and levels of its nucleotide exchange factors HSPBP1 and HSP70E (Bracher and Verghese, 2015) increased 19.6- and 1.4-fold, respectively, at the 48 h time point. A divergent behavior between chaperone and co-chaperones was also observed for HSP90A: the abundance of HSP90A increased in the *70B*-amiR line 1.6-fold 48 h after the N source shift as did its co-chaperones AHA1, HOP1, SGTA1, CYN40, and FKB62 (cochap 1, on average by 1.7-fold), whereas the abundance of its co-chaperones TPR2, CCPP23, CCPP5, and CNS1 decreased on average by 40% (cochap 2) (Schroda and deVitry, 2023). The eight subunits of the TRiC/CCT chaperonin, involved in the folding of specific substrates including actin and tubulin (Yam et al., 2008), declined on average by 50% in the *70B*-amiRNA versus control line. While proteasome subunits increased mildly by 20% on average, proteins involved in autophagy showed a 5.6-to 55-fold increased abundance in the *70B*-amiRNA versus control line at the 48 h time point (Figure 4F).

The response of ER-localized chaperones was also quite distinct, with Hsp70 homolog BIP2 and nucleotide exchange factor HSP70G increasing 10- and 1.4-fold in abundance, respectively, in the *70B*-amiRNA versus control line 48 h after N shift, while BIP1, HSP90B, and calreticulin CRT2 (Perez-Martin et al., 2014) declined mildly and calnexin CNX1 strongly (Figure 4G). Mitochondrial HSP70C and its co-chaperone MGE1 as well as chaperonin CPN60C and its co-chaperonin CPN10 increased mildly by up to 1.4-fold in the *70B*-amiRNA versus control line at the 48 h time point, while AAA+ ATPase chaperones CLPX1, HSLU1 and HSLU2 as well as the proteolytic subunit HSLV1 (Schroda and deVitry, 2023) increased between 1.6- and 14-fold (Figure 4H).

Proteins of the 60S and 40S subunits of cytosolic ribosomes declined on average by up to 53% at the 48 h and 72 h time points in the *70B*-amiRNA versus control line (Figure 4I). In contrast. chloroplast ribosomal subunits showed peculiar kinetics, with proteins of both subunits increasing mildly in *70B*-amiR by 10-15% 24 h after the N shift. The abundance of proteins of the 50S subunit then declined on average by 25% and 50% at the 48 h and 72 h time points, respectively, while proteins of the 30S subunit remain elevated at the 48 h time point and declined by 25% at the 72 h time point.

### HSP70B depletion results in a decline of proteins involved in energy conversion in chloroplasts and mitochondria and major metabolic pathways

The abundances of proteins forming the four major complexes of the photosynthetic light reactions – PSII, PSI, cytochrome b_6_/f, PSI, ATP synthase as well as the antennae of both photosystems – continuously declined in the *70B*-amiR line to about 50% of the abundance of the control line at the 48 h and 72 h time points (Figure 4J). The only exceptions among the overall 57 quantified proteins were PETO, which declined to 10% of control line levels, and LHCBM9 and LHCB7, which increased 26- and 5-fold, respectively. PETO is involved in cyclic electron flow (Takahashi et al., 2016), while LHCBM9 and LHCB7 have proven and proposed roles in photoprotection, respectively (Sawchuk et al., 2008; Grewe et al., 2014). Strikingly, the abundance of proteins forming the four major complexes of the mitochondrial OXPHOS system – complexes I to V – also showed a continuous decline in the *70B*-amiR versus control line (Figure 4K). While proteins of complexes I, II, III and V declined on average to about 50% of control line levels at time point 48 h and 72 h, those of complex IV declined even to less than 10%.

Finally, proteins of the Calvin-Benson cycle (CBC), the TCA cycle and gluconeogenesis declined in the *70B*-amiR line at time points 48 h and 72 h to about 50%, 40% and 14% of control line levels, respectively (Figure 4L). The only exception among the overall 39 enzymes monitored was CBC enzyme ribulose-5-phosphate 3-epimerase (RPE1), pointing to another role of this enzyme beyond the CBC.

Overall, the proteomics data reveal dramatic proteome remodeling upon HSP70B depletion within and beyond the chloroplast compartment.

### HSP70B depletion results in formation of aberrant structures at thylakoid conversion zones

To monitor potential changes in thylakoid ultrastructure, we took transmission electron micrographs of control and *70B*-amiR lines before and after shifting the N-source (Figure 5A, Supplemental Figure S4A). Thylakoid membranes were indistinguishable between control and *70B*-amiR cells grown with ammonium as N source. However, already 24 h after the N source shift we observed aberrant structures at sites where multiple thylakoid membranes converge, specifically in the *70B*-amiR lines. Such aberrant structures have been observed previously in VIPP1 depleted cells and in cells exposed to heat stress for more than 12 h and were referred to as prolamellar body (PLB)-like structures (Nordhues et al., 2012; Hemme et al., 2014). Serial sections at such sites revealed several distinct white areas of variable size but unknown chemical composition (Figure 5B, Supplemental Figure S4B).

**Figure 5.**
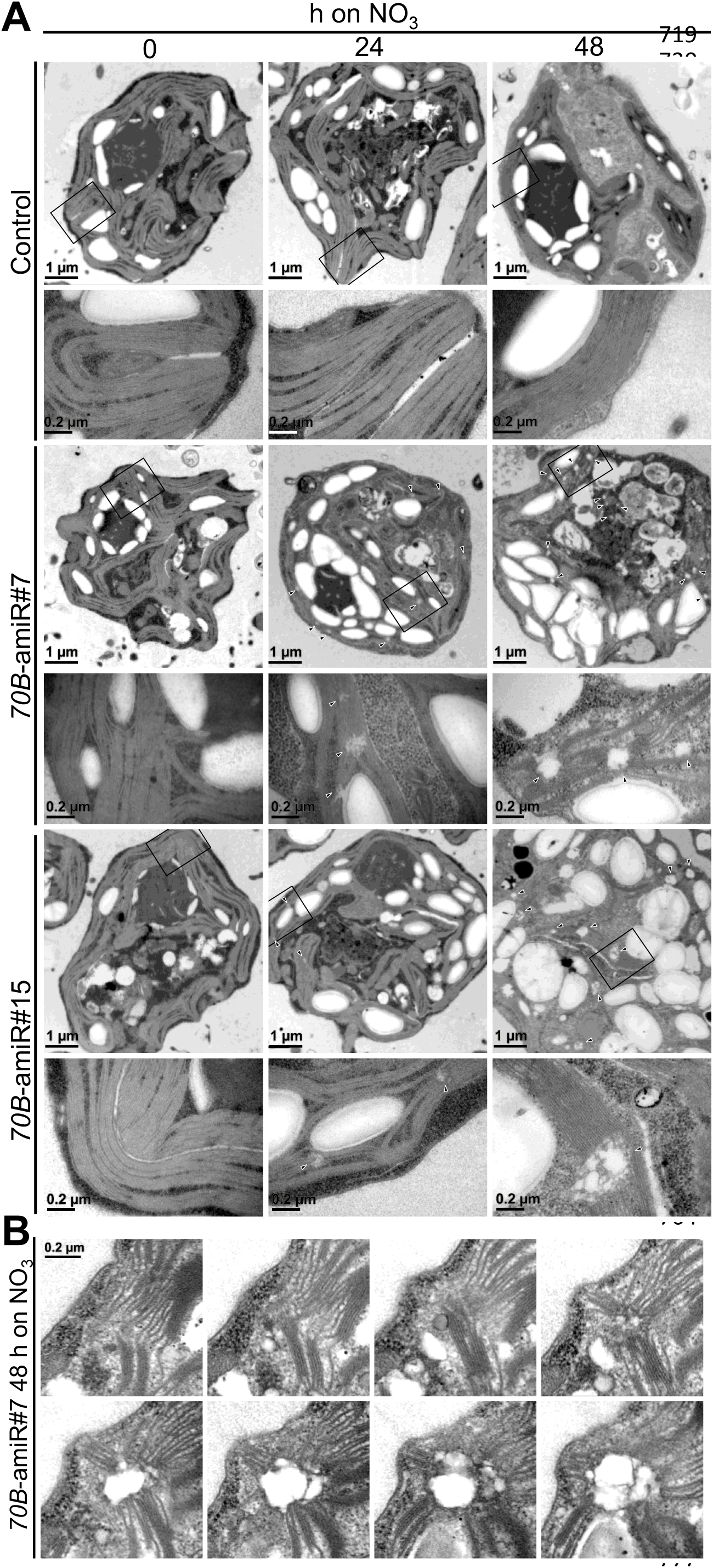
Documentation of PLB-like structures in *70B*-amiR lines. **A)** Electron microscopy images of cells from control and two *70B*-amiR lines #7 and #15 before (0 h) and 24 h and 48 h after shifting the N source from ammonium to nitrate. Cells were grown at 60 µmol photons m^-2^ s^-1^. For each line and time point, an overview image is shown on the top and zoom-ins of the regions demarcated by black boxes are shown on the bottom. Triangles indicate PLB-like structures at thylakoid conversion zones. Bars in overview images = 1 µm, those in zoom-ins = 0.2 µm. **B)** Serial sections of a PLB-like structure in a cell of the *70B*-amiR line #7 48 h after shifting to nitrate as N source. The scale bar (0.2 µm) holds for all images. Additional images and serial sections are shown in Supplemental Figure S4.

The electron micrographs also revealed a larger size of *70B*-amiR cells especially at the 48 h time point, in line with the increased cell diameter measured by the Coulter Counter (Figure 1D). We also noticed an increased accumulation of starch grains especially in the 48 h *70B*-amiR cells (Figure 5A, Supplemental Figure S4A).

### VIPP1 shifts into higher molecular weight assemblies upon depletion of HSP70B

HSP70B together with its co-chaperones CDJ2 and CGE1 was shown to catalyze the assembly of VIPP1 monomers/dimers to oligomers as well as the disassembly of VIPP1 oligomers in vitro (Liu et al., 2007). Similarly, a function of Arabidopsis chloroplast cpHsc70 in VIPP1 oligomer dynamics has recently been demonstrated (Li et al., 2025). We therefore wanted to test whether the depletion of HSP70B affected the VIPP1 assembly state in vivo. To this end, we shifted WT and *70B*-amiR cells from ammonium to nitrate medium for 36 h and fractionated cells taken during the time course into soluble and insoluble fractions by freeze-thaw cycles followed by centrifugation. In WT cells, only a small portion of VIPP1 was recovered in the insoluble pellet fraction during the entire time course (Figure 6A). This was observed also for VIPP1 in *70B*-amiR cells until 12 h after the N source shift. After 24 h, however, about one quarter and after 36 h more than half of VIPP1 shifted into the insoluble pellet fraction. We wondered whether the increased partitioning of VIPP1 to the insoluble fraction was due to increased membrane binding or due to increased oligomerization. To test this, we repeated the N source shift, solubilized WT and *70B*-amiR cells harvested during the time course with 2% Triton X-100, and precipitated insoluble proteins by an ultracentrifugation at 435,000 g for 1 h. As shown in Figure 6B, the fraction of pelleted VIPP1 did not change during the time course in WT cells, while it roughly doubled in *70B*-amiR cells 12 h after the N-shift and dramatically increased 24 h after the shift. To get an idea about the oligomeric nature of VIPP1, we centrifuged Triton X-100 solubilized WT and *70B*-amiR cells harvested after the N source shift through a 10-30% sucrose gradient (Figure 6C). VIPP1 from WT cells remained to the very large part on top of the gradient, while only a small portion migrated into the first gradient fraction and an even smaller portion migrated through the gradient. These three portions likely correspond to VIPP1 monomers/dimers, small and very large assembly states, respectively, as observed previously (Liu et al., 2007). VIPP1 from *70B*-amiR cells showed the same migration pattern before the N shift, but at 12 h and later time points after the N shift was increasingly found in larger and very large assemblies. At the same time, the amount of VIPP1 in the solubilized membranes on top of the gradient decreased. Taken together, these data indicate an important role of HSP70B in sustaining VIPP1 oligomer dynamics.

**Figure 6.**
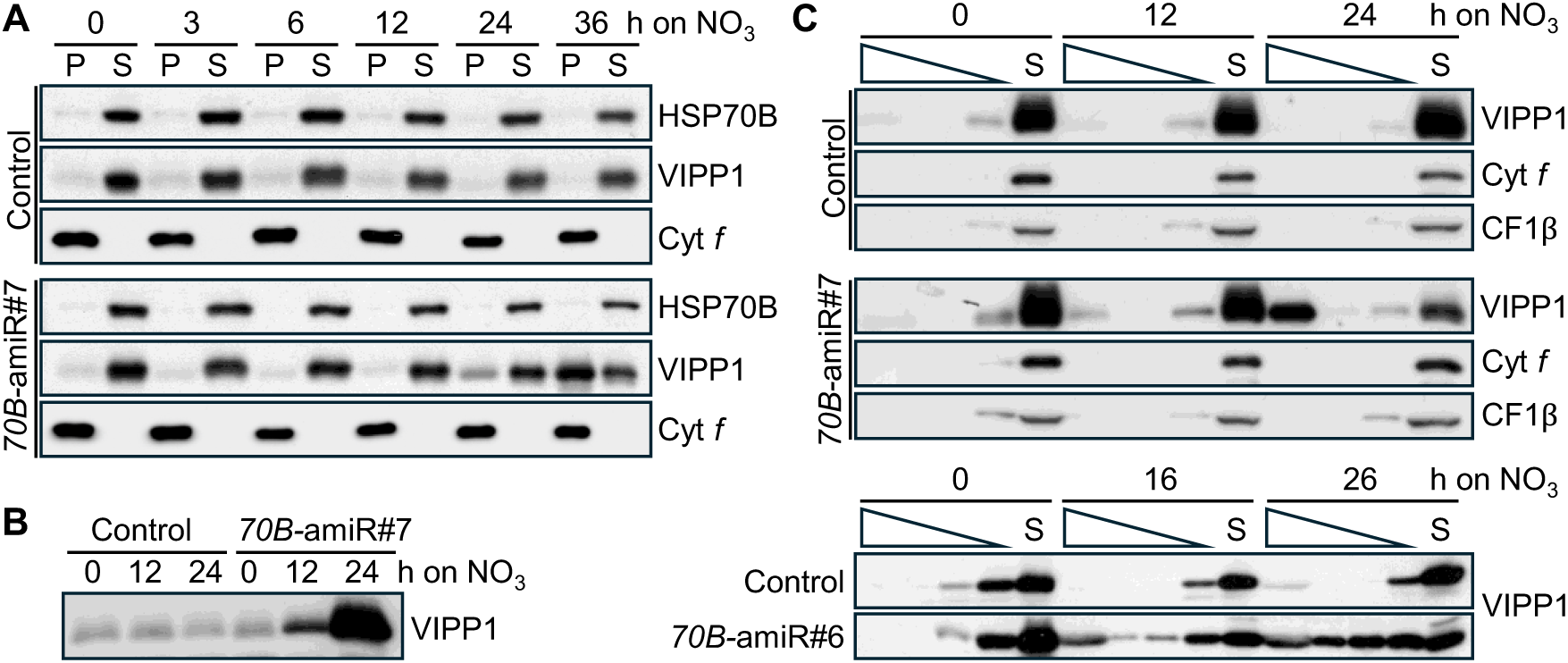
Monitoring the VIPP1 oligomeric state upon HSP70B depletion. **A)** Analysis of VIPP1 solubility. Control and *70B*-amiR line #7 harvested before and after switching the N source for the indicated time points were lysed by freeze-thaw cycles and separated into soluble (S) and insoluble (P) fractions by a 35-min centrifugation at 16,100 *g*. Pelleted proteins were resuspended in the same volume of buffer as used for cell lysis to allow for quantitative comparison and analysed by immunoblotting. HSP70B and Cyt *f* served as markers for soluble and membrane-integral proteins, respectively. **B)** Analysis of Triton-insoluble protein complexes. Total proteins were extracted in the presence of 2% Triton X-100 from control and *70B*-amiR line #7 at the indicated time points after the N source shift. Proteins in very large complexes were then pelleted by a 1-h centrifugation at 435,000 *g* and analysed for the presence of VIPP1 by immunoblotting. **C)** Analysis of high molecular weight oligomers of VIPP1. Cells of control and *70B-*amiRNA lines #7 (top) and #6 (bottom) were harvested before and after switching the N source at the indicated time points and solubilized with 2% Triton X-100. Lysates were subjected to a 2-h centrifugation at 79,000 *g* on 10-30% sucrose gradients and the soluble material on top of the gradient (S) and gradient fractions were analysed by immunoblotting.

### Cells depleted of HSP70B are high light sensitive

The mild reduction of HSP70B levels by the expression of an *HSP70B*-antisense RNA resulted in an increased sensitivity of the antisense line to high light intensities (Schroda et al., 1999). This phenotype was observed only with one antisense line and only at very high light intensities. Having several *HSP70B*-amiRNA lines in two different strain backgrounds at hand, we set out to validate this phenotype. For this, we shifted cultures of *70B*-amiR cells from ammonium to nitrate medium and grew them at low light intensities for 24 h. Cultures were subsequently exposed to high light intensities of 1000 µmol photons m^-2^ s^-1^ for another 24 h. As shown in Figure 7 and Supplemental Figure S5, two different *70B*-amiR lines in both UVM4-NIT and cw15-325 cell backgrounds bleached after 12 and 16 h high light exposure, respectively, while no bleaching was observed in the controls. High light exposure resulted in a drop in Fv/Fm values from ∼0.75 to ∼0.24 in control and *70B*-amiR line @22. However, while the control line recovered Fv/Fm to 0.48 during high light exposure, no recovery was observed for the *70B*-amiR line (Figure 7B). Hence, we could reproduce the high light sensitive phenotype observed in the *HSP70B*-antisense line.

**Figure 7.**
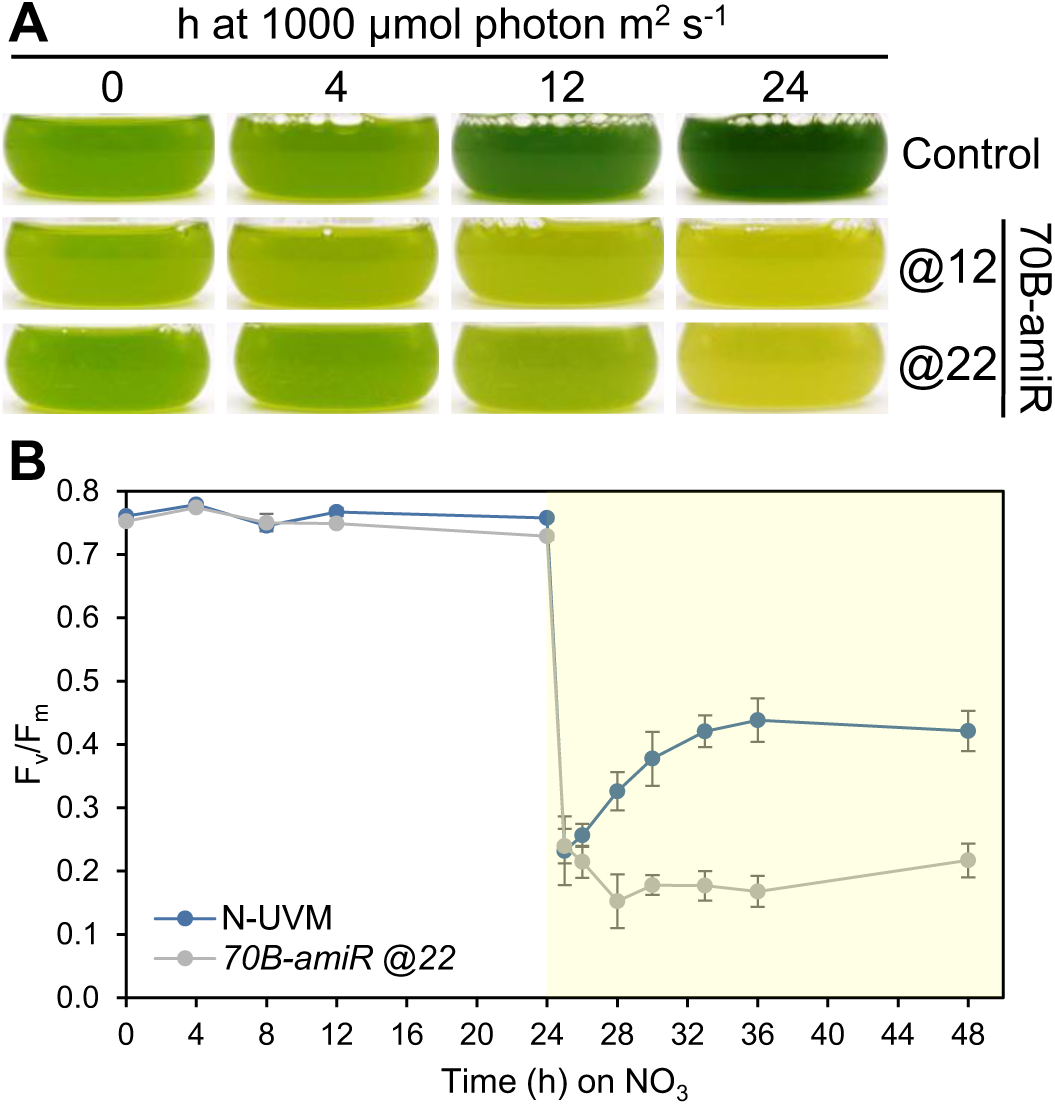
Testing the sensitivity of *70B*-amiR lines to high light intensities. **A)** Cultures of UVM4-NIT (control) and *70B*-amiR lines @12 and @22 were shifted from ammonium to nitrate medium and cultivated for 24 h at 60 µmol photons m^-2^ s^-1^ to deplete HSP70B (t = 0). Cultures were then exposed for 24 h to 1,000 µmol photons m^-2^ s^-1^. Cultures were diluted 1:4 with fresh nitrate-containing medium after 12 h of high light treatment. **B)** Monitoring of Fv/Fm values in control and *70B*-amiR line @22 in the experiment depicted in A). The yellow surface indicates the high light period. Shown are mean values from three biological replicates, error bars depict SDs.

## Discussion

### HSP70B affects VIPP1 oligomer dynamics

HSP70B depletion resulted in the accumulation of VIPP1 in complexes of higher molecular weight (Figure 6). In vitro, VIPP1 was shown to spontaneously assemble into baskets of stacked rings (Gupta et al., 2021; Liu et al., 2021), helical rods (Liu et al., 2007; Theis et al., 2019), and flat spirals/polygons/carpets on membranes (Junglas et al., 2025; Naskar et al., 2025; Pan et al., 2025). However, of these structures, only helical rods have been observed in situ by cryo-ET (Gupta et al., 2021; Gachie et al., 2025). These rods tubulate membranes, and some appear to connect the inner envelope with thylakoids, presumably to facilitate lipid transfer (Gupta et al., 2021).

Since VIPP1 forms these oligomeric structures in vitro without energy input, it is conceivable that an ATP-consuming machinery is required for their disassembly in vivo. Eukaryotic and archaeal ESCRT-III systems achieve this via the AAA+ ATPase Vps4 (Babst et al., 1998; Mierzwa et al., 2017), for which there are no homologs in bacterial ESCRT-III systems, including plastids (Williams and Low, 2025). We have previously shown that HSP70B, together with its co-chaperones CDJ2 and CGE1, can disassemble VIPP1 rods into smaller assembly states in vitro (Liu et al., 2007). Moreover, in Arabidopsis, chloroplast Hsc70-1 was shown to play a similar role in regulating VIPP1 dynamics (Li et al., 2025). These findings suggest that the HSP70/DnaK system may fulfil, in prokaryotic systems, a function analogous to that of Vps4 in eukaryotes in mediating ESCRT-III oligomer disassembly.

We cannot say which of the oligomeric structures formed by VIPP1 accumulate upon HSP70B depletion in vivo (Figure 6). The nature of larger VIPP1 oligomers formed in the absence of cpHsc70-1 also remained unresolved in Arabidopsis (Li et al., 2025). However, we observed lesions in thylakoid conversion zones upon depletion of HSP70B (Figure 5) as well as upon depletion of VIPP1 (Nordhues et al., 2012). This suggests that VIPP1 function is impaired not only when its abundance is reduced, but also when its oligomerization dynamics are perturbed, despite an overall increase in VIPP1 abundance and additional expression of VIPP2. This interpretation is supported by the albino phenotype shared by Arabidopsis *cpHsc70*-amiRNA lines and *vipp1* knockout lines (Latijnhouwers et al., 2010; Zhang et al., 2012).

The ability of HSP70B to maintain VIPP1 oligomer dynamics may also become limiting under prolonged heat stress, when HSP70B is largely engaged in assisting protein folding. This could explain the formation of lesions in thylakoid conversion zones under such conditions (Hemme et al., 2014), as well as the aggravation of phenotypes in *cphsc70-1* knockout lines upon heat stress (Su and Li, 2008; Li et al., 2025).

### Why is HSP70B essential?

We show that inducible depletion of chloroplast HSP70B to less than 30% of WT levels resulted in cell division arrest, increased cell size, and accumulation of starch granules (Figures 1, 5A; Supplemental Figures S2–S4). Starch accumulation in Chlamydomonas cells is frequently observed following nutrient starvation, including nitrogen, sulfur, phosphate, potassium, and magnesium depletion (Ball et al., 1990), but also in cells exposed to prolonged heat stress (Hemme et al., 2014). The essential nature of chloroplast Hsp70s is conserved, as knocking out genes encoding stromal HSP70 (Hsp70-2) in the moss *Physcomitrium patens* (Shi and Theg, 2010) and simultaneously those encoding cpHsc70-1 and cpHsc70-2 in Arabidopsis is lethal (Su and Li, 2008)

How can HSP70B depletion result in cell division arrest? For the following reasons, we believe that this phenotype is not primarily related to HSP70B’s role in maintaining VIPP1 dynamics.

First, the function of HSP70B on the VIPP1 oligomeric state is considered to be mediated by the co-chaperone CDJ2. Accordingly, CDJ2 was upregulated 9.7-fold 48 h after induction of *HSP70B*-amiRNA expression (Figure 4A), most likely as a compensatory response. Importantly, a Chlamydomonas *cdj2* mutant is viable (Kafri et al., 2023), suggesting that impaired regulation of VIPP1 by HSP70B is unlikely to explain the cell division arrest.

Second, *VIPP1*-RNAi and -amiRNA lines exhibit the same aberrant structures in thylakoid conversion zones as observed in *70B*-amiR lines. Both grow normally under low light intensities, but are sensitive to high light (Nordhues et al., 2012) (Figure 7; Supplemental Figure S5). Thus, perturbation of VIPP1 function alone does not result in growth arrest under low light.

Hence, the essential function of HSP70B in the chloroplast is most likely mediated by another of the six J-domain proteins in the Chlamydomonas chloroplast (Chiu et al., 2013; Trösch et al., 2015). In the Chlamydomonas Library project (CLiP) (https://phytozome-next.jgi.doe.gov/), no insertional mutants exist for *HSP70B* and *CDJ1*, and of the eight mutants in *CDJ5* seven have the insertion in the 3′-UTR and one in an intron. In contrast, several insertion mutants in the coding sequence exist for *CDJ3*, *CDJ4*, and *CDJ6*. This distribution suggests that the essential role of HSP70B may be linked to its function in protein folding mediated by CDJ1. This could affect the folding of individual labile proteins such as deoxyxylulose 5-phosphate synthase (DXS), the key enzyme of the MEP pathway, which was shown to be a cpHsc70 substrate in Arabidopsis (Pulido et al., 2016). Impaired synthesis of key metabolites through such pathways could mimic nutrient depletion, resulting in cell division arrest, starch accumulation, and induction of autophagy (Ball et al., 1990; Perez-Perez et al., 2010). Alternatively, depletion of HSP70B may more generally impair the folding of newly synthesized or imported proteins, resulting in protein aggregation. These aggregates would then trigger the chloroplast unfolded protein response (cpUPR), as evidenced by the drastic upregulation of the typical cpUPR marker proteins HSP22E/F/C, CLPB3, and DEG1C (Figure 4BC) (Gabelmann and Schroda, 2025), as well as autophagy markers (Figure 4F). This response resembles that observed upon depletion of the chloroplast ClpP protease (Ramundo et al., 2014). Whether VIPP2, another typical marker of the cpUPR, is upregulated to compensate for loss of VIPP1 function (Nordhues et al., 2012) or as part of the cpUPR remains unclear.

Alternatively, the essential nature of HSP70B could also be related to a function mediated by CDJ5 which, dependent on the presence of its 4Fe–4S cluster, was found in proximity to proteins related to the regulation of photosynthetic electron transfer and photosystem biogenesis (König et al., 2025). The loss of CDJ5 functionality could explain the decline of virtually all components of the photosynthetic light reactions upon HSP70B depletion (Figure 4J).

Upon HSP70B depletion, none of the highly upregulated chloroplast-targeted proteins, including VIPP1, VIPP2, DEG1C, CLPB3, CDJ1, or HSP22E/F, accumulated as precursors (Figure 2), as would be expected if import were impaired (Jarvis et al., 1998). This argues against a major role of HSP70B in chloroplast protein import, as previously suggested for chloroplast Hsp70s in moss and Arabidopsis (Shi and Theg, 2010; Su and Li, 2010), and is consistent with recent structures of the import machinery indicating that the Ycf2–FtsHi complex constitutes the chloroplast protein import motor (Jin et al., 2022; Liu et al., 2023; Liang et al., 2024).

The dramatic remodeling of the proteomes in and beyond the chloroplast compartment upon HSP70B depletion (Figure 4) makes it difficult to distinguish processes affected by impaired HSP70B function from processes resulting from cell division arrest. In future work, it will therefore be important to analyse and compare the proteomes in cells lacking individual chloroplast J-domain proteins.

### The sensitivity of HSP70B-depleted cells to high light intensities is most likely related to its role in maintaining VIPP1 dynamics

The high light sensitivity of cells expressing an *HSP70B* antisense construct was previously suggested to be due to a possible role of HSP70B in facilitating PSII repair (Schroda et al., 1999; Schroda et al., 2001; Yokthongwattana et al., 2001). Since the high light-sensitive phenotype was also observed in *VIPP1*-RNAi and *VIPP1*-amiRNA lines exhibiting similar aberrant structures at thylakoid conversion zones as *70B*-amiR lines (Nordhues et al., 2012), we propose that the phenotype is more likely related to the role of HSP70B in maintaining VIPP1 oligomer dynamics rather than to a direct role in PSII repair.

The molecular mechanisms underlying high light sensitivity in cells impaired in VIPP1 functionality remain unclear. Nevertheless, the upregulation upon HSP70B depletion of LHCBM9 and LHCB7, with proven and proposed roles in photoprotection, respectively (Sawchuk et al., 2008; Grewe et al., 2014), is noteworthy (Figure 4J). Since this upregulation occurred in *70B*-amiR cells grown at low light intensities, it may indicate impaired electron flow from the light reactions to downstream electron sinks.

A large-scale screen for *Chlamydomonas* mutants with impaired photosynthesis identified 115 genes whose disruption caused a growth defect at 750 µmol photons m⁻² s⁻¹ under photoautotrophic conditions, but not under heterotrophic conditions in the dark (Kafri et al., 2023). *CDJ2* was among these genes, suggesting that the *cdj2* mutant may display high-light sensitivity, similar to *VIPP1*- and *HSP70B*-amiRNA lines. A detailed characterization of the *cdj2* mutant could therefore help elucidate the consequences of impaired VIPP1 oligomer disassembly by the chloroplast HSP70B machinery, while avoiding the pleiotropic effects observed in *70B*-amiR lines that arise from the multiple and essential functions of HSP70B.

## Methods

### Vector construction

The amiRNA targeting Chlamydomonas *HSP70B* was designed using the <u>W</u>eb <u>M</u>icroRNA <u>D</u>esigner (WMD3) webtool at http://wmd3.weigelworld.org (Ossowski et al., 2008) following the instructions for Chlamydomonas given by (Molnar et al., 2009). The resulting oligonucleotide amiFor-70B and amiRev-70 (Supplemental Table S1)) were annealed by boiling at 100°C and slow cooling down to 60°C (0.3°C/min) in a thermocycler (Biometra). After purification, the annealed oligos were ligated into *Spe*I-digested pMS539 (Schmollinger et al., 2010), yielding pMS542. To substitute *ARG*7 with the *aadA* cassette as a selection marker, a PCR was performed on pMBS542 with primers HSP70B-ami-for and VIPP1-ami-rev (Supplemental Table S1) using the Q5 high-fidelity DNA polymerase (M0491, New England Biolabs) according to the manufacturer’s protocol. This resulted in a 1510-bp PCR product comprising the *NIT1* promoter, the *HSP70B*-amiRNA and the *RPL12* terminator flanked by *Bbs*I recognition sites and position 2F fusion sites defined by the MoClo system (Weber et al., 2011). The resulting PCR-product was combined with pCM1-01 (*PSAD* promotor + *aadA* gene + *PSAD* terminator) (Crozet et al., 2018) and end-linker pICH41744 and cloned into the level 2 destination vector pAGM4673 (Weber et al., 2011) by simultaneous digestion with *Bbs*I and ligation with T4 DNA ligase yielding pMBS822.

### Strains and culture conditions

*Chlamydomonas reinhardtii* strain cw15-325 (*cwd mt+ arg7*), kindly provided by R. Matagne (University of Liège, Belgium), was used as recipient strain for transformation with control constructs pCB412 (Schroda et al., 1999) and pMS539 (Schmollinger et al., 2010) or *70B-*amiRNA construct pMS542. All three plasmids contain the wild-type *ARG7* gene and were linearized by digestion with *Hind*III prior to transformation. The UVM4 strain (Neupert et al., 2009) was equipped with the wild-type *NIT1* and *NIT2* genes to generate UVM4-NIT previously (Probst et al., 2025). Plasmid pMBS888 was linearized with *Eco*RV prior to transformation. Transformation into cw15-325 and UVM4-NIT was done by agitation of 1µg linear DNA with glass beads (Kindle, 1990). Strains were grown mixotrophically in TAP medium (Kropat et al., 2011) containing 7.5 mM ammonium, referred to as TAP-NH_4_, on a rotatory shaker at 25°C at 60 µE m^-2^ s^-1^. To exchange the nitrogen source, cells cultivated on TAP-NH_4_ medium to a maximal cell density of 5 x 10^6^ cells/mL were centrifuged at 4,000 *g* for 2 min at 25°C. The resulting cell pellet was resuspended in TAP medium containing 7.5 mM nitrate (TAP-NO_3_) to wash out the remaining ammonium and pelleted. Finally, the cell pellet was resuspended in TAP-NO_3_ medium. If necessary, cells were diluted during the experiment with fresh TAP-NO_3_ medium to ensure that cell densities never exceeded 5 x 10^6^ cells/mL and cells remain in log phase.

### Cell number and cell size distribution

Cell number and cell size distribution were determined using a Coulter-Counter Z2 (Beckman-Coulter). Each measurement was carried out in two technical replicates and the average was used for further comparisons. The size distribution was monitored by counting cells in 256 windows between 4.4 µm and 12.2 µm. For comparison between different strains in different cell densities, the data was normalized to the highest number of cells in a single window, which was set to 1 (% max).

### Immunoblot analyses

Protein extraction, quantification, SDS-PAGE using 10% polyacrylamide gels, and immunoblotting using semidry blotting via a discontinuous transfer system and enhanced chemiluminescence (ECL) was performed as described previously (Laemmli, 1970; Liu et al., 2005). Antisera described previously targeting HSP70B (Schroda et al., 1999), CDJ1 (Willmund et al., 2008), VIPP1 (Liu et al., 2005), VIPP2 (Nordhues et al., 2012), HSP90C (Willmund and Schroda, 2005), CLPB3 (Kreis et al., 2023), DEG1C (Theis et al., 2019), HSP22E/F (Rütgers et al., 2017), HSP90A (Schulz-Raffelt et al., 2007), CF1β (Lemaire and Wollman, 1989), Cyt *f* (Pierre and Popot, 1993), and ATG8 (Perez-Perez et al., 2010). The antiserum against PsaA and the anti-rabbit IgG–HRP secondary antibody were from Agrisera (AS06 172 and AS09 602, respectively).

### Sucrose density centrifugation

Sucrose density centrifugation to separate high molecular weight oligomers of VIPP1 from smaller assemblies was performed as described previously (Liu et al., 2007). Briefly, 2 x 10^8^ cells were harvested and resuspendend in 1 mL KH buffer (20 mM Hepes–KOH pH 7.2, 80 mM KCl) containing 2% Triton X-100 and 0.25 x protease inhibitor cocktail (Roche). Cells were lysed by sonication on ice and centrifuged for 2 min at 16,100 *g*, 4°C in a table centrifuge to remove non-solubilized matter. The supernatant was transferred on top of a continuous 10 mL 10-30% sucrose gradient and centrifuged for 2 hours at 79,000 *g*, 4°C.

### High light treatment

High light (1000 µmol photons m^-2^ s^-1^) treatment was done with cultures in beakers placed onto a water-cooled metal plate on a rotary shaker to keep the temperature constant at 25°C. For lines in the cw15-325 strain background, 50-mL cultures in 150-mL beakers were illuminated by Osram HLX 250W 64663 Xenophot bulbs, with a glass plate nonpermissive to infrared and UV irradiation placed between light source and beakers. For lines in the UVM4-NIT strain background, high-light treatment was performed with 45-mL cultures in 100-mL Erlenmeyer flasks using the light system CXB3590-X4, COB 4×100W (CF Grow).

### Whole-cell proteomics

For whole-cell proteome analysis, cells corresponding to 20 µg total protein were precipitated with four volumes of cold acetone (80% final concentration) at −20 °C for 3 h and pelleted by centrifugation (25,000 *g*, 15 min, 4 °C). Pellets were washed with 80% acetone, briefly air-dried, and resuspended in 20 µL 8 M urea, 25 mM NH_4_HCO_3_. Proteins were reduced and alkylated by adding 2 µL of freshly prepared 0.2 M TCEP (tris(2-carboxyethyl)phosphine), 0.8 M chloroacetamide, followed by a 20-min incubation at 25°C in the dark. Samples were then diluted with 25 mM NH_4_HCO_3_ to a final urea concentration of 4 M, Lys-C was added at a ratio of 1:100 (w/w, Lys-C/protein) and digestion allowed to take place for 3 h at 37° C. Samples were further diluted with 25 mM NH4HCO3 to a final urea concentration of 1 M and supplemented with acetonitrile to a final concentration of 5%. Trypsin was then added at a ratio of 1:75 (w/w, trypsin/protein) and proteins allowed to digest overnight at 37° C. To complete digestion, more trypsin was added to yield a ratio of 1:40 and the samples were incubated for another 3 h at 37° C. Digestion was stopped by adding formic acid and trifluoroacetic acid at final concentrations of 2% and 0.2%, respectively. Peptides were desalted using C18 StageTips, eluted with 60% acetonitrile containing 0.1% trifluoroacetic acid and 1% formic acid, dried, and subjected to LC–MS/MS analysis using a nanoElute UHPLC system coupled to a timsTOF Pro2 mass spectrometer (Bruker Daltonics) according to (John et al., 2024).

Raw data were processed with FragPipe v19.1 and searched against the *Chlamydomonas reinhardtii* genome (release v5.6) (Merchant et al., 2007; Craig et al., 2022) including organellar proteins. The resulting protein intensities were analyzed in Perseus (v1.6.15.0) (Tyanova et al., 2016). Filtering for contaminants, log2 transformation, normalization and imputation of the values was performed with the same parameters used in (Kreis et al., 2023). Proteins that were not identified in at least three out of four replicates were removed. Differentially abundant proteins were identified by two-sample t-tests and considered significant at q < 0.05 with |log2 fold change| ≥ 1. Proteins with log2 fold change ≥ 1 were classified as upregulated and those with log2 fold change ≤ −1 as downregulated. The evaluated MS data have been deposited at https://git.nfdi4plants.org/probsta/HSP70B_amiRNA_Chlamy.git. The principal component analysis was performed based on the processed data using five components, Benjamini-Hochberg as cutoff method with a FDR of 0.05 and relative enrichment based on ‘protein’ as selected parameter.

### Transmission electron microscopy

Embedding for electron microscopy was performed according to a protocol published previously (Nordhues et al., 2012). 10^7^ cells were harvested and fixed for 2 hours in 2.5% glutaraldehyde buffered with 100 mM Na-cacodylate (pH 7.2). After washing, cells were fixed for 1 hour in 1% OsO_4_ at 4°C. After incubating the cells for 15 min in 20% BSA (w/v), the cell pellet was fixed for another 30 min with 2.5% glutaraldehyde. The cell pellet was embedded in 1.5% agarose and then cut into several pieces (∼1 to 2 mm^3^ in size), dehydrated in ethanol and embedded in glycid ether 100 (Serva) with propylene oxide as intermediate solvent. Ultrathin sections (60 to 70 nm) were cut with a diamond knife (type ultra 35°, Diatome) on an EM UC6 ultramicrotome (Leica) and mounted on single-slot Pioloform-coated copper grids (Plano). The sections were stained with uranyl acetate and lead citrate (Reynolds, 1963) and viewed with a JEM-2100 (Jeol) or EM 902A (Carl Zeiss) transmission electron microscope (both operated at 80 kV). Micrographs were taken using a 4080 x 4080-pixel or 1350 x 1040-pixel charge-coupled device camera (UltraScan 4000 or Erlangshen ES500W, respectively; Gatan) and Gatan Digital Micrograph software (version 1.70.16). Image brightness and contrast were adjusted and figures assembled using Adobe Photoshop 8.0.1.

## Supplemental Files

**Supplemental Figure S1.** Schematic diagram of the vector pMS542 for the inducible expression of an amiRNA against *HSP70B*.

**Supplemental Figure S2.** Growth analysis and HSP70B protein content of all *70B*-amiR lines used in this study.

**Supplemental Figure S3.** Growth analysis of *70B*-amiR lines in the UVM4-NIT strain background.

**Supplemental Figure S4.** Documentation of aberrant structures at the origin of multiple thylakoid membranes in *70B*-amiR lines.

**Supplemental Figure S5.** Testing the sensitivity of *70B*-amiR lines to high light intensities.

**Supplemental Table S1.** Primers used for amiRNA constructs.

**Supplemental Data Set S1.** Shotgun proteomics data comparing relative abundances of proteins between 70B-amiR line @12 and the UVM4-NIT control.

## Acknowledgments

We thank Olivier Vallon for the antisera against CF1β and Cyt *f*. This work was supported by the Deutsche Forschungsgemeinschaft [RTG 2737 and SFB/TRR175, project C02].

## Author Contributions

A.P. and S.S. generated the constructs and transgenic lines and performed all experiments supported by J.B. and D.S. A.-K.U. and S.G. performed the TEM analysis. F.S ran the mass spectrometry analyses. M.S. conceived and supervised the project and wrote the article with contributions from all authors.

