## Supplemental Data for "Depletion of Chloroplast HSP70B Triggers Proteostasis Collapse and Compromises Thylakoid Membrane Integrity in *Chlamydomonas*"

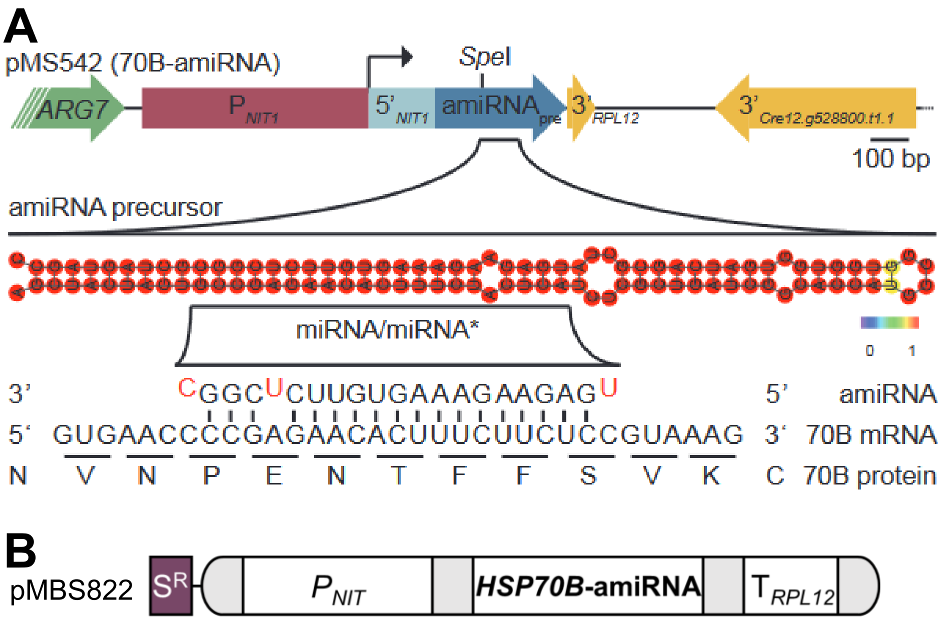


**Supplemental Figure S1.** Schematic diagram of the vector pMS542 for the inducible expression of an amiRNA against *HSP70B*. **A)** pMS542 contains the *ARG7* gene as a selection marker (green). The nitrate reductase promoter ($P_{NIT1}$, red) with its 5'-UTR (light blue) controls the expression of the cre-MIR1157 miRNA precursor (dark blue) (Molnar et al., 2009), which contains an amiRNA directed against *HSP70B* in its unique SpeI cleavage site. The 3'-UTR of the *RPL12* gene (yellow) serves as a transcriptional terminator. The secondary structure of the produced amiRNA precursor was predicted using RNAfold on the Vienna Webserver (Gruber et al., 2008). The sequence of the amiRNA, the target region in the *HSP70B* mRNA and the deduced amino acid sequence at the target region are shown below. **B)** pMBS822 contains the *aadA* cassette conferring resistance against spectinomycin ($S^R$), the *NIT1* promoter driving expression of cre-MIR1157 containing the same amiRNA as depicted in pMS542, and the *RPL12* terminator.
Supports Figure 1.

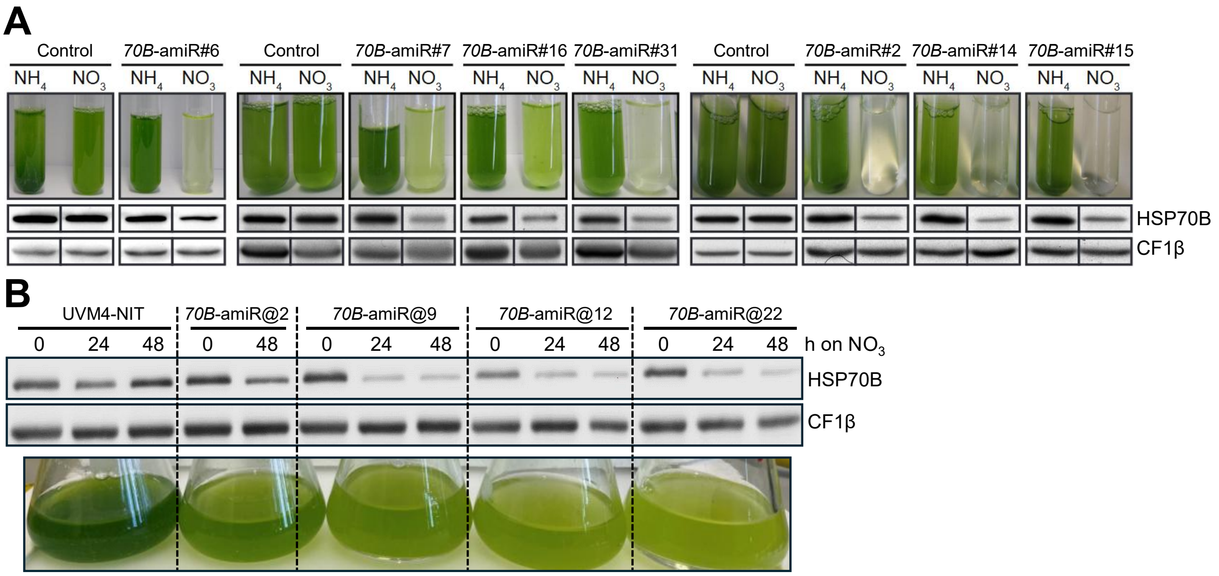


**Supplemental Figure S2.** Growth analysis and HSP70B protein content of all *70B*-amiR lines used in this study. **A)** *70B*-amiR lines in the cw15-325 strain background (#). Control and *70B*-amiR lines were inoculated in TAP medium containing ammonium ($NH_4$) or nitrate ($NO_3$) at 2 x $10^4$ cells/mL and grown for 3 days, after which the pictures were taken. For protein analyses, cultures were grown to 5 x $10^6$ cells/mL in TAP-$NH_4$ when the medium was changed to TAP-$NO_3$. Total cell protein extracts from samples taken before and 24 h after switching the N source corresponding to 0.5 µg chlorophyll were then analyzed by immunoblotting to compare contents of HSP70B relative to the loading control CF1β. **B)** *70B*-amiR lines in the UVM4-NIT strain background (@). Control and *70B*-amiR lines were grown to stationary phase in TAP-$NH_4$, diluted 1:10 with TAP-$NH_4$ and grown for 16 h. Then cells were transferred to TAP-$NO_3$ and samples were taken after 0 h, 24 h and 48 h. Cultures were diluted again 1:4 with TAP-$NO_3$ 16 h before the 48-h sample was taken. Pictures were taken 48 h after the medium change. Total cell protein extracts corresponding to 0.5 µg chlorophyll were then analyzed by immunoblotting to compare contents of HSP70B relative to the loading control CF1β.
Supports Figure 1.

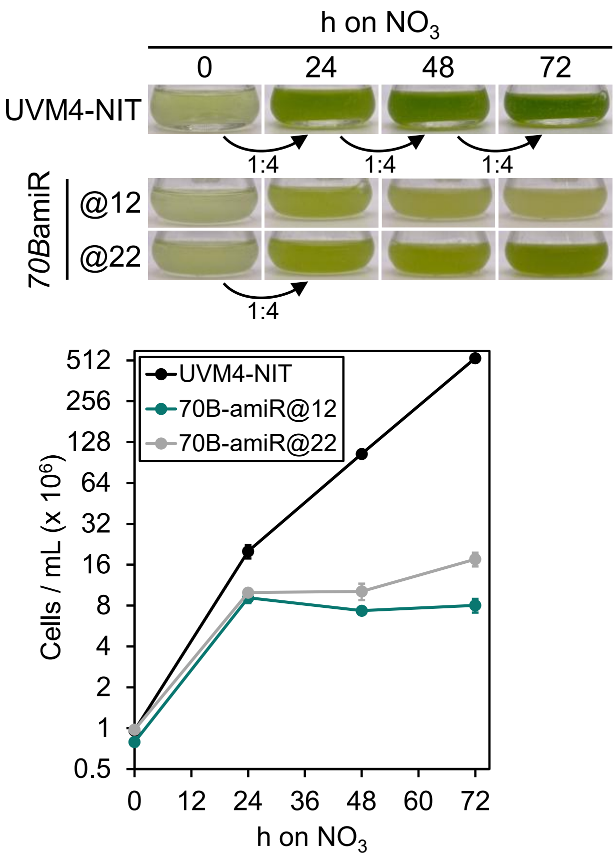


**Supplemental Figure S3.** Growth analysis of *70B*-amiR lines in the UVM4-NIT strain background. Cells of the UVM4-NIT recipient strain and two lines expressing the *70B*-amiR construct pMBS888 (Supplemental Figure S1B) were grown to log-phase in TAP-$NH_4$. Cells were then diluted to $10^6$ cells/mL in TAP-$NO_3$. Cell growth was monitored during 72 h with photographs of the cultures taken (top) and cell densities measured (bottom). Cultures were diluted 4-fold with TAP-$NO_3$ every 24 h. Error bars represent standard deviation (n = 3). Supports Figure 1.

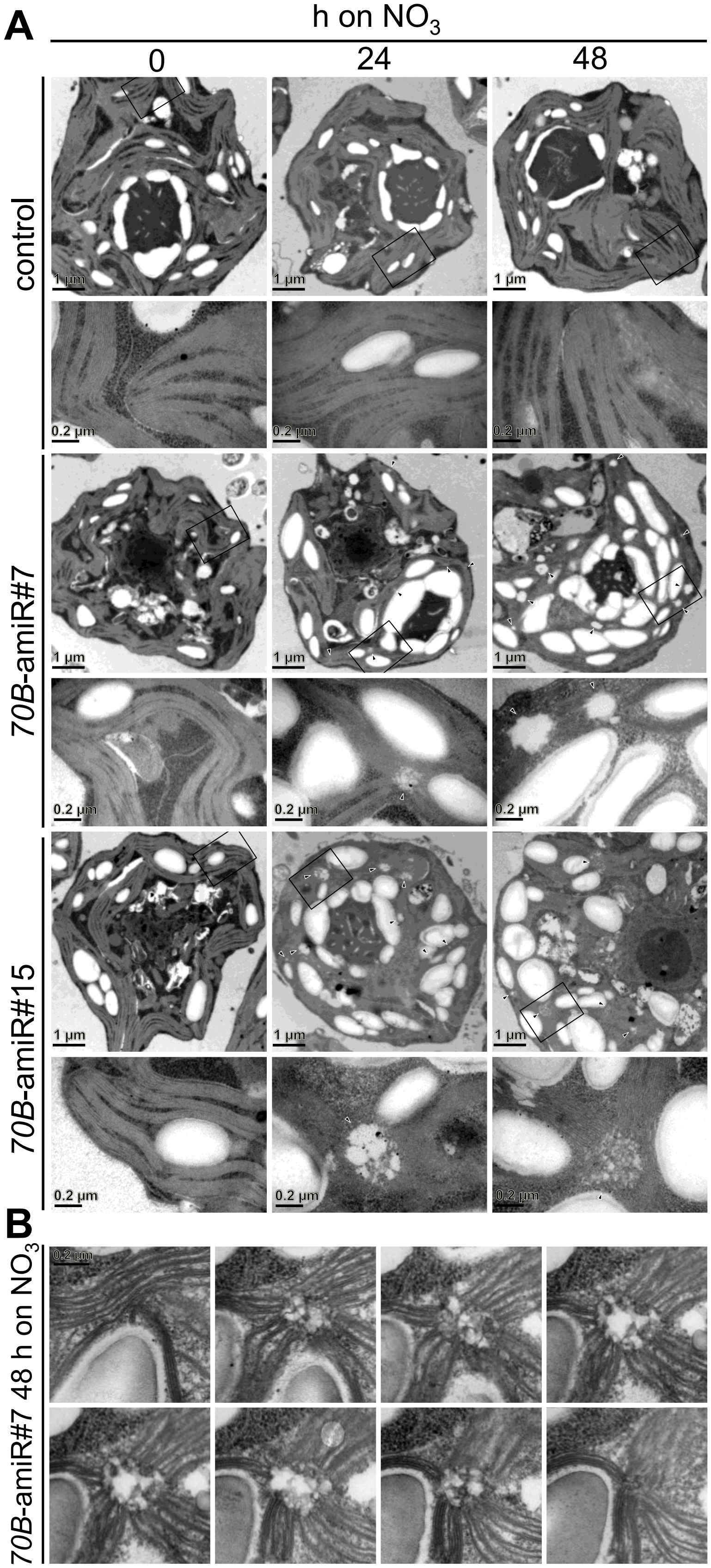


**Supplemental Figure S4.** Documentation of PLB-like structures in *70B*-amiR lines. **A)** Electron microscopy images of cells from control and two *70B*-amiR lines #7 and #15 before (0 h) and 24 h and 48 h after shifting the N source from ammonium to nitrate. Cells were grown at 60 µmol photons $m^{-2}$ $s^{-1}$. For each line and time point, an overview image is shown on the top and zoom-ins of the regions demarcated by black boxes are shown on the bottom. Triangles indicate PLB-like structures at thylakoid conversion zones. Bars in overview images = 1 µm, those in zoom-ins = 0.2 µm. **B)** Serial sections of a PLB-like structure in a cell of the *70B*-amiR line #7 48 h after shifting to nitrate as N source. The scale bar (0.2 µm) holds for all images. Supports Figure 5.

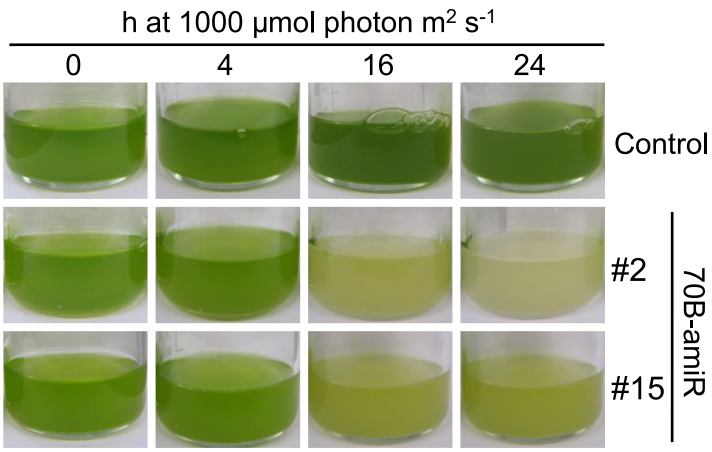


**Supplemental Figure S5.** Testing the sensitivity of *70B*-amiR lines to high light intensities. Cultures of Control and *70B*-amiR lines #2 and #15 were shifted from ammonium to nitrate medium and cultivated for 24 h at 60 µmol photons $m^{-2}$ $s^{-1}$ to deplete HSP70B (t = 0). Cultures were then exposed for 24 h to 1,000 µmol photons $m^{-2}$ $s^{-1}$. Cultures were diluted 1:4 with fresh nitrate-containing medium after 16 h of high light treatment.

**Supplemental Table S1.** Primers used for amiRNA constructs. Uppercase letters indicate sequences targeting the *HSP70B* mRNA and sequences in the pMS542 vector, respectively.

| Primer | Sequence | Target |
|---|---|---|
| amiFor-70B | 5'-ctagtGCCGAGAACACTTTCTACTCAtctcgctgatcggcacca tggggtggtggtgatcagcgctaTGAGAAGAAAGTGTTCTCGGCg-3' | HSP70B |
| amiRev-70B | 5'-ctagcGCCGAGAACACTTTCTTCTCAtagcgctgatcaccacc acccccatggtgccgatcagcgagaTGAGTAGAAAGTGTTCTCGGCa-3' | |
| HSP70B-ami-for | 5'-ttgaagacaagcAAGGTGACCCGCCAGCCCCCGCTCCT-3' | pMS542 |
| VIPP1-ami-rev | 5'-ttgaagacaataGTTCGCCGTCAACGTGCCATGGATAC-3' | |
